# Insights into intraspecific diversity of central carbon metabolites in *Saccharomyces cerevisiae* during wine fermentation

**DOI:** 10.1101/2023.02.24.529865

**Authors:** Ludovic Monnin, Thibault Nidelet, Jessica Noble, Virginie Galeote

## Abstract

*Saccharomyces cerevisiae* is a major actor in winemaking that converts sugars from the grape must into ethanol and CO_2_ with outstanding efficiency. Primary metabolites produced during fermentation have a great importance in wine. While ethanol content contributes to the overall profile, other metabolites like glycerol, succinate, etc., also have significant impacts, even when present in lower concentrations. *S. cerevisiae* is known for its great genetic diversity that is related to its natural or technological environment. However, its range of metabolic diversity which can be exploited to enhance wine quality depends on the pathway considered. Our experiment assessed the diversity of primary metabolites production in a set of 51 *S. cerevisiae* strains from various genetic backgrounds. Results pointed out great yield differences depending on the metabolite considered, with ethanol having the least variation. A negative correlation between ethanol and glycerol was observed, confirming glycerol synthesis as a suitable lever to reduce ethanol yield. Genetic groups were linked to specific metabolic yields such as α-ketoglutarate or acetate. This research highlights the potential of using natural yeast diversity in winemaking and provides detailed data set on metabolite production of well known (ethanol, glycerol, acetate) or little-known (lactate) primary metabolites production.

## 1. Introduction

Fermented products have today a great importance in human societies, both economically and socially. Throughout history, humans and fermentation have shared a long path: the first trace of cereal fermentation has been found in Israel and estimated to date back to 13000 B.C. (Liu *et al*., 2018) and the first known fermented beverage from rice, honey, and a fruit, has been traced back to 7000 B.C. in China (McGovern *et al*., 2004). Since then, fermentation uses have expanded into a wide diversity of processes and products, such as food, beverages or more recently biofuels. In alcoholic beverages, alcoholic fermentation is the main step of elaboration and is mostly carried out by yeasts of the *Saccharomyces* genus, especially the species *Saccharomyces cerevisiae*. A perfect example is wine, which is the result of the alcoholic fermentation of grapes or grape juice. From a technological point of view, wine fermentation is the biotransformation of glucose and fructose, existing in equal proportions in grapes, into carbon dioxide and ethanol, which imparts new characteristics to the product: sensory qualities, stability…

Alcoholic fermentation is of high technological interest as well as of metabolic importance for *Saccharomyces cerevisiae*. Through glycolysis, this biological process generates pyruvate and energy in the form of ATP. Pyruvate, which is a central metabolite, is then converted in two steps into ethanol and carbon dioxide, which ensures a rapid reoxidation of enzymatic cofactors used in glycolysis, making alcoholic fermentation the most efficient way to promptly supply energy to the cell (Bakker *et al*., 2001). Moreover, in typical wine making conditions, this is the only way for *S. cerevisiae* to produce ATP, respiration being repressed by the Crabtree effect or impossible due to the absence of dioxygen (De Deken, 1966; Pfeiffer and Morlay, 2014). The main products of fermentation, carbon dioxide and ethanol, are by far the most produced metabolites during alcoholic fermentation and therefore in wine making (Nidelet *et al*., 2016). A simple way to compare these productions between species, strains or fermentation conditions is to define yield as the ratio of the quantity of metabolite produced per unit of substrate consumed. Ethanol yield in wine fermentations carried out by *S. cerevisiae* is known to reach around 0.47 gram per gram of hexoses consumed, which represent 92% of the maximum theoretical yield (calculated as one mole of glucose producing two moles of ethanol) (Tilloy *et al*., 2015). Most of the remaining hexoses are used as a carbon source for cell multiplication and the production of other metabolites in minor concentrations, such as glycerol, acetate, succinate, acetaldehyde, lactate, etc. These metabolites represent much lower carbon fluxes, but can be of significant technological value. Glycerol, which is linked to stress resistance, can impact wine mouthfeel above a certain concentration (Albertyn *et al*., 1994; Noble and Bursick 1984). It has been identified as the second most produced metabolite in fermentation and as the flux with the greatest impact on ethanol production (Goold *et al*., 2017). Acetate, which is a way to restore redox balance in the cell and a metabolic intermediary, is a major off-flavour-linked compound and subject to legal limits (Vilela-Moura *et al*., 2008). It appears that yields of fermentation metabolites such as ethanol, glycerol or acetate are linked to the degree of domestication of the producing strains (Tapia *et al*., 2018).

In most studies, the carbon metabolites considered are those most present in fermentation, which are final steps of metabolic pathways and therefore important markers: ethanol, glycerol, succinate, α-ketoglutarate… However, other carbon metabolites are evoking an increasing interest, as they can deeply shape the sensorial identity of wine, in particular malic acid (Vion *et al*., 2023) or lactic acid (with a particular focus on yeast species other than *S. cerevisiae*, which is commonly considered as a very poor producer) (Vicente et *al*., 2021).

For all compounds, yield values differ according to strain and fermentation conditions (oxygenation, temperature, nutrients concentrations, presence of other microorganisms…) (Du *et al*., 2012; Tronchoni *et al*., 2022) but the range of variation for ethanol remains very limited compared to that observed for biomass or other metabolites. In their work, Nidelet *et al*. (2016) compared 43 strains from six different ecological origins and showed that the coefficient of variation of carbon flux toward ethanol synthesis following glycolysis and alcoholic fermentation is only between 2 and 3%. In a contrasting way, yields of glycerol or acetate have a respective variation of around 10 and 30% although they represent significantly lower carbon fluxes for the cell (Camarasa *et al*., 2011; Nidelet *et al*., 2016). Generally, global yields are calculated at fixed points of the fermentation: at 80% of hexoses consumed, during the exponential phase, etc… One of the reasons for these choices is that ethanol yield is not constant during fermentation so that the flux is difficult to calculate, apart during the exponential growth phase which is the only stage with a quasi-steady state (Celton *et al*., 2012; Nidelet *et al*., 2016; Quirós *et al*., 2013). However, in a wine production context, a yield per strain can only be calculated when fermentation is completed, i.e. when all hexoses have been consumed.

While constituting a minor fraction of the cell carbon flux, other metabolites are synthesised at very low concentrations. Despite their low levels, these metabolites have a significant impact on the final fermented product, with examples including organic acids, higher alcohols and esters.. Consequently, their production mechanisms have been extensively studied (Antonelli *et al*., 1999; Regodón Mateos *et al*., 2006).

Over the past thirty years, considerable research efforts have been made to understand and influence primary metabolism, mainly with the aim of reducing wine final ethanol content. Beside physical or chemical methods, many microbial strategies have been developed to modify ethanol production during fermentation. We can cite here genetically modified yeast strains, hybrids strains, optimisation through adaptive laboratory evolution… (reviewed in Varela and Varela, 2019). However, modulating the central carbon metabolism (CCM) without disturbing the cell balance still remains complex in a wine context, mostly because of the multigenic basis of the associated traits (Bro *et al*., 2006; Hubmann, Foulquié-Moreno, *et al*., 2013; Hubmann, Mathé, *et al*., 2013; Salinas *et al*., 2012). Nevertheless, elaborating approaches to develop *S. cerevisiae* strains with a modified primary metabolic yield in wine fermentation requires to clearly identify the diversity of central metabolism as well as its constraints and trade-offs. In this context, our study presents the outcomes of a screening strategy applied to 51 strains from different origins in order to identify the range of variability in primary fermentation metabolite yields under laboratory wine-like conditions among the species *S. cerevisiae*.

## 2. Materials and methods

### 2.1 Strains

51 strains of *S. cerevisiae* were used (see information in supplementary data (S1)). Strains were selected considering results from precedent works in the laboratory, with the aim to maximise diversity in fermentation profiles (Camarasa *et al*., 2011; Legras *et al*., 2018; Nidelet *et al*., 2016). EC1118 was chosen as a reference strain to estimate block effect. Genetically modified (GM) and laboratory strains evolved for precise CCM traits were also included. Strains were conserved at −80 °C in 20% glycerol YPD medium (10 g/L yeast extract, 20 g/L peptone, 20 g/L glucose) and cultivated on YPD agar plate (YPD + 20 g/L agar).

### 2.2 Genetic groups constitution

Strains from various genetic backgrounds, but all linked to fermented beverages, are represented in the set. 37 out of 51 of them have had their genome sequenced in previous works (Akao *et al*., 2011; Eder *et al*., 2018; Fay and Benavides, 2005; Liti *et al*., 2009; Marsit *et al*., 2015; Novo *et al*., 2009; Schacherer *et al*., 2009). To classify and organise intraspecific diversity, two works were used to define the following genetic groups: wine, rum, West African, sake and flor (Legras *et al*., 2018; Peter *et al*., 2018). Strains without information were labelled as ‘Unknown’. A supplementary group, labelled as ‘Miscellaneous’, was used to assemble those strains with mosaic, very singular or unclassifiable genomes, but was not used as a consistent group like the others.

### 2.3 Fermentation conditions

Fermentation conditions were chosen to ensure a quick and complete alcoholic fermentation. One colony was grown overnight in YPD medium as pre-culture. Then 10^6^ cells/ml of this pre-culture were inoculated in 280 ml fermenters. A synthetic medium that mimics grape must was used for fermentation, based on the work of Bely *et al*. (1990). This medium contained, per litre: 90 g glucose, 90 g fructose, 6 g citric acid, 6 g DL-malic acid, 750 mg KH_2_PO_4_, 500 mg K_2_SO_4_, 250 mg MgSO_4_.7H_2_O, 155 mg CaCl_2_.2H_2_O, 200 mg NaCl, 4 mg MnSO_4_.H_2_O, 4 mg ZnSO_4_.7H_2_O, 1 mg CuSO_4_.5H_2_O, 1 mg KI, 0.4 mg CoCl_2_.6H_2_O, 1 mg H_3_BO_3_, 1 mg (NH_4_)_6_Mo_7_O_24_, 20 mg myo-inositol, 2 mg nicotinic acid, 1.5 mg calcium pantothenate, 0.25 mg thiamine-HCl, 0.25 mg pyridoxine and 0.003 mg biotin. 425 mg/L of yeast assimilable nitrogen (YAN) was added as a mixture of amino acids and ammonium containing, per litre: 460 mg NH_4_Cl, 612 mg L-proline, 505 mg L-glutamine, 179 mg L-tryptophan, 145 mg L-alanine, 120 mg L-glutamic acid, 78 mg L-serine, 76 mg L-threonine, 48 mg L-leucine, 45 mg L-aspartic acid, 45 mg L-valine, 38 mg L-phenylalanine, 374 mg L-arginine, 33 mg L-histidine, 33 mg L-isoleucine, 31 mg L-methionine, 18 mg L-glycine, 17 mg L-lysine, 18 mg L-tyrosine and 13 mg L-cysteine. Fermenters filled with medium were heat-sterilised (100°C, 10 min) and cooled down to 28°C before inoculation.

Fermentations were carried out at 28°C with agitation. Fermenter weight was measured twice a day to observe fermentation progress, weight loss being directly a consequence of carbon dioxide production and release. Fermentations carried at the same time represented a fermentation block. Three replicates were made for each strain (except LMD17, LMD37, LMD39, performed in six replicates due to their use in a parallel project and EC1118 performed in duplicate per block, *i. e*. 28 replicates in total).

### 2.4 Metabolite analysis

Fermentation metabolites concentrations were measured using high performance liquid chromatography (HPLC) as described in Deroite *et al*. (2018) and analysing chromatograms with OPEN LAB 2X software. Fermentation samples were centrifuged 5 min at 3500 rpm at 4 °C and kept at -18°C. Before analysis, samples were diluted to 1/6 with 0.0025 mol/L H_2_SO_4_ and then centrifuged 5 min at 13000 rpm at 4 °C. The supernatant was kept at -18°C until analysis. The HPLC method allowed the measurement of concentrations of glucose, fructose, ethanol, glycerol, acetate, succinate, α-KG, lactate and malate. Analyses were performed in duplicate and the mean was calculated for each sample and used in results analysis.

Quantification was made with a Rezex ROA column (Phenomenex, Torrance, California, USA) set at 60 °C on an HPLC equipment (HPLC 1260 Infinity, Agilent Technologies, Santa Clara, California, USA). It was resolved isocratically with 0.0025 mol/L H_2_SO4 at a flow rate of 0.6 mL/min. Concentrations of acetate, malate, lactate and α-KG were measured with a UVmeter at 210 nm and other compounds with a refractive index detector.

For each fermentation, two measurements were taken: in the must before fermentation (for each block) and at the end of fermentation. All analyses were conducted on finished fermentation, i.e. when combined fructose and glucose concentration fell below 3 g/L, or when fermenter weight stayed constant for 24h.

Yield was calculated for each metabolite as follows:

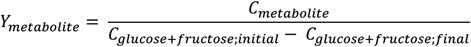

Concentrations were expressed in g/L or mg/L, leading to yields expressed in g/g or mg/g. When necessary, yield values were expressed as follows: mean ± standard error.

Malate being both produced and consumed during fermentation, the final concentration measured was used to calculate a difference with the initial theoretical malate concentration (here 6 g/l), the initial concentration not having been measured.

### 2.5 Statistical analysis

Statistical analysis was performed using R studio software (version: 1.4.1106). Multiple R packages were used to carry out data analysis and visualisation: ‘tidyverse’ (1.3.2), ‘FactoMineR’ (2.6), ‘factoextra’ (1.0.7), ‘ggpubr’ (0.4.0), ‘viridis’ (0.6.2) ‘GGally’ (2.1.2)

EC1118 was used in each fermentation block in order to evaluate a possible block effect (i.e. differences between reference values from different fermentation blocks). It was estimated using the following model:

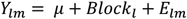

With: Y the phenotype (yield for a given metabolite) for the block *l* (1-51) and the replicate *m* (1-2). *μ* represent the mean of the considered phenotype and *E* the residual error, with *E ∼ N*(0, σ^2^).

A block effect was observed on EC1118 data. This was corrected by calculating a variation factor on EC1118 metabolite values. This correction (raw value - correction coefficient) was then applied to all variables. The block effect being eliminated, yields can be expressed with the following model:

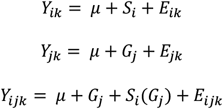

With: Y the phenotype (yield for a given metabolite) corrected for block effect for the strain *i* (1-51), the genetic group *j* (1-5) and for the replicate *k* (1-28). *μ* represent the mean of the considered phenotype, *S* the effect of the strain *i, G* the effect of the genetic group j and *E* the residual error, with *E ∼ N*(0, σ^2^). To express the yield variation for a metabolite among a group of strains, the variation coefficient was used (Albatineh *et al*., 2014). A correction according to the number of strains in a group was applied, allowing us to compare groups of different sizes. The correction was applied as follows:

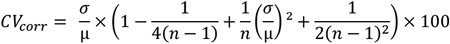

With, for a group of strains and a metabolic yield: *μ* the mean, *σ* the standard deviation, *n* the size of the group and *CV*_*corr*_ the corrected coefficient of variation, expressed as percentage. The coefficient of variation was calculated for ethanol, glycerol, acetate, succinate, α-KG and lactate yields (malate being excluded).

### 2.6. Comparison with other screening works

For the four main metabolites considered (ethanol, glycerol, acetate and succinate), we compared the results of the present work with significant datasets previously obtained. Data were acquired online or directly from the authors. These data compared different sets of strains in similar conditions but with different analysis timing (resumed in Table 1). To overcome these differences, a normalised yield value was calculated as follows:

**Table 1.**
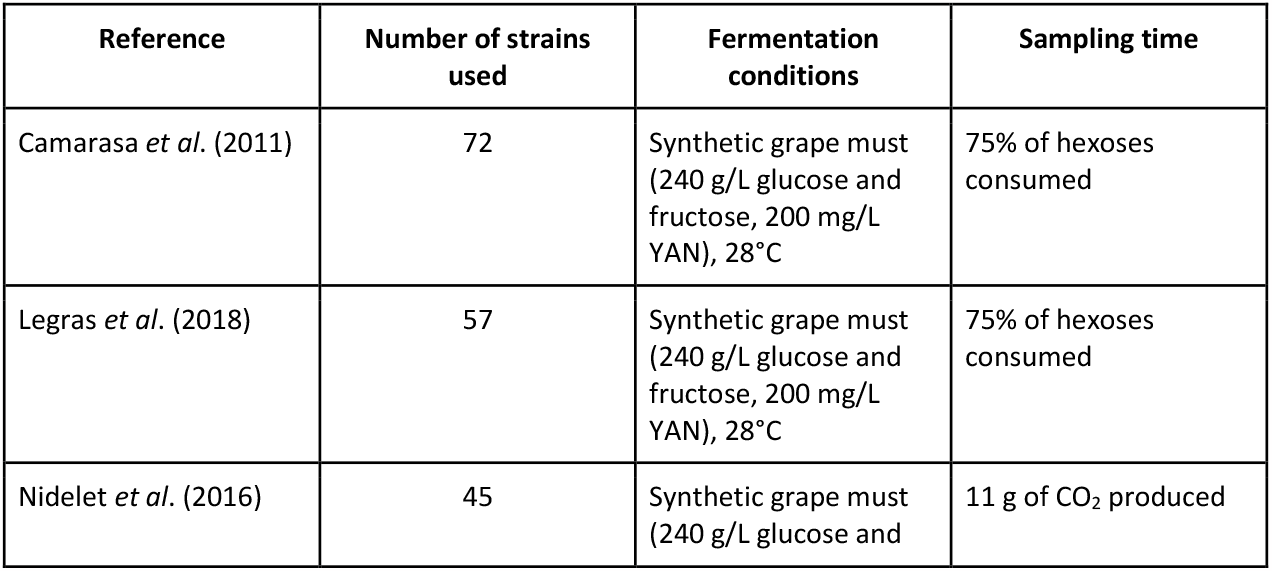
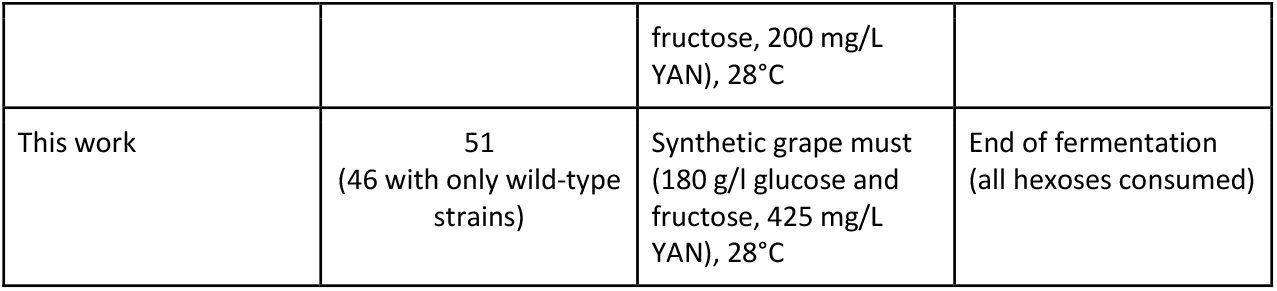
List of screening works on CCM in fermentation used as comparison.

| Reference | Number of strains used | Fermentation conditions | Sampling time |
| --- | --- | --- | --- |
| Camarasa *et al.* (2011) | 72 | Synthetic grape must (240 g/L glucose and fructose, 200 mg/L YAN), 28°C | 75% of hexoses consumed |
| Légras *et al.* (2018) | 57 | Synthetic grape must (240 g/L glucose and fructose, 200 mg/L YAN), 28°C | 75% of hexoses consumed |
| Nidelet *et al.* (2016) | 45 | Synthetic grape must (240 g/L glucose and fructose, 200 mg/L<br>YAN), 28°C | 11 g of CO<sub>2</sub> produced |
| This work | 51<br>(46 with only wild-type<br>strains) | Synthetic grape must<br>(180 g/l glucose and<br>fructose, 425 mg/L<br>YAN), 28°C | End of fermentation<br>(all hexoses consumed) |

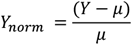

With: *Y* the yield of a metabolite for a designated strain, *μ* the mean of all strains and *Y*_*nom*_ the normalised yield. For the data set produced in the present work, normalisation was performed with and without optimised strains.

## 3. Results

Here we present results obtained for 51 strains following the fermentation of a synthetic grape must. Concentrations were measured for 7 compounds linked to the central carbon metabolism (CCM) : ethanol, glycerol, succinate, acetate, α-KG, lactate and malate, determined by HPLC. After correction of the block effect (see Material & Method), strains were compared to each other. Among our set of strains, two are genetically modified: LMD41 and LMD45 (confidential genetic constructions) and three have been obtained using adaptive laboratory evolution methods: 5074, LMD13, LMD14 (Cadière *et al*., 2011; Tilloy *et al*., 2014). All these strategies aimed to modify CCM during fermentation. These strains were used as controls for their metabolic productions. Consequently, for correlation and PCA, they were withdrawn because their metabolism does not represent a natural variation within the species. The 46 other strains were placed in a group termed ‘wild-type’. All strains were able to entirely consume glucose and fructose in the must within 5 days (data not shown). In addition to the overall analysis, separate results for each metabolite are available as supplementary information.

### 3.1 Comparison of variation of metabolic yields

In the aim to have a better comparison of metabolic yields between them and between strain groups, the coefficient of variation (CV) was calculated for each metabolite (Figure 1). Malate is a component of the synthetic medium used and is both produced and consumed during fermentation. Moreover, its final delta outcomes can be positive or negative depending on strains; consequently, the application of CV as a descriptive parameter for malate becomes impractical due to its versatile behaviour in our experimental context.

**Figure 1.**
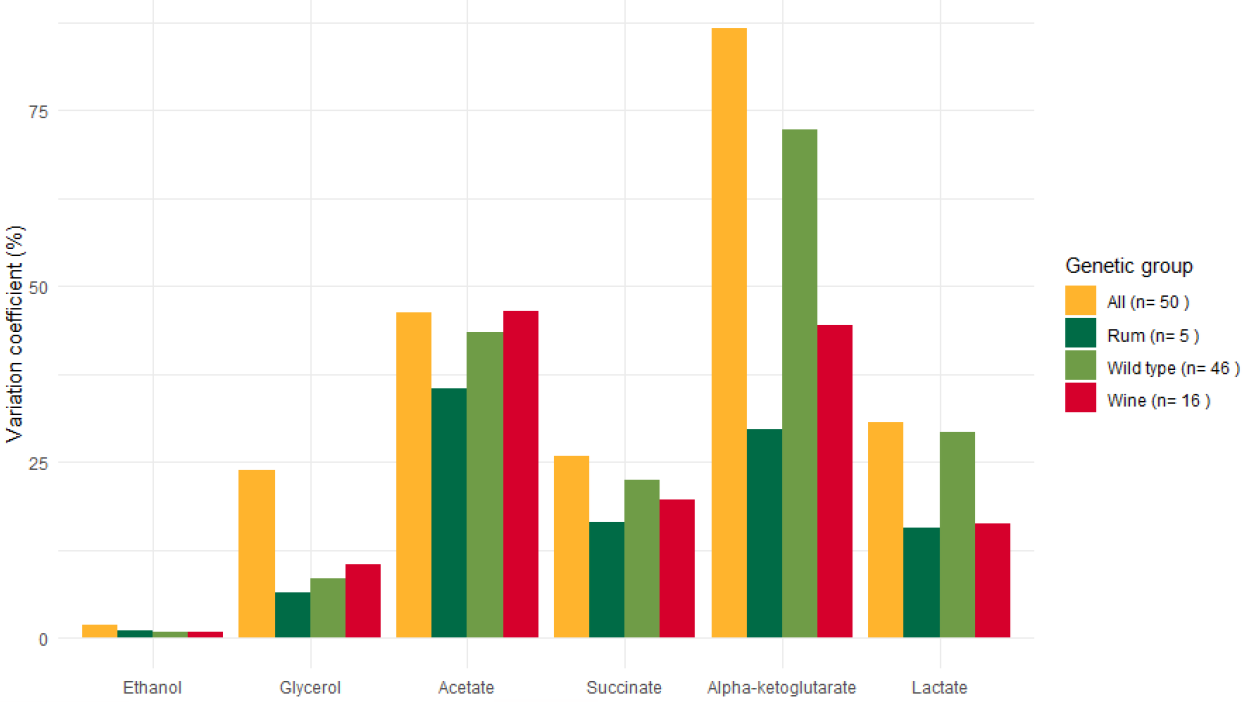
Variation coefficient of each metabolite for all strains (n = 51), wild-type strains (n = 46), wine strains (n = 16) and rum strains (n = 5)

The CV was corrected to allow comparison of groups of different sizes (described in Material & Method). Among all metabolites examined, ethanol presented the lowest yield variation with a variation coefficient of 1.8 % when all strains were considered. When only wild-type strains were taken into account, the coefficient of variation was even lower, dropping to 0.8 %. The wine and rum groups exhibited similar values. Other metabolites displayed a more important variation among our selection of strains, with a peak for α-KG around 86% (when all strains are considered). Overall, variation was higher when considering all strains. Acetate was the only exception, with a similar coefficient of variation both for the whole set and wine group (both ∼46%).

### 3.2 Correlations between metabolic yields

Pearson’s coefficients were used to compare all metabolites with one another and evaluate possible correlations. The malate concentration used corresponds to the difference between its initial concentration and the concentration observed at the end of fermentation. Results, represented as a correlogram, are available in Figure 2 and 3, respectively for the whole set of strains and for wild-type strains only.

**Figure 2.**
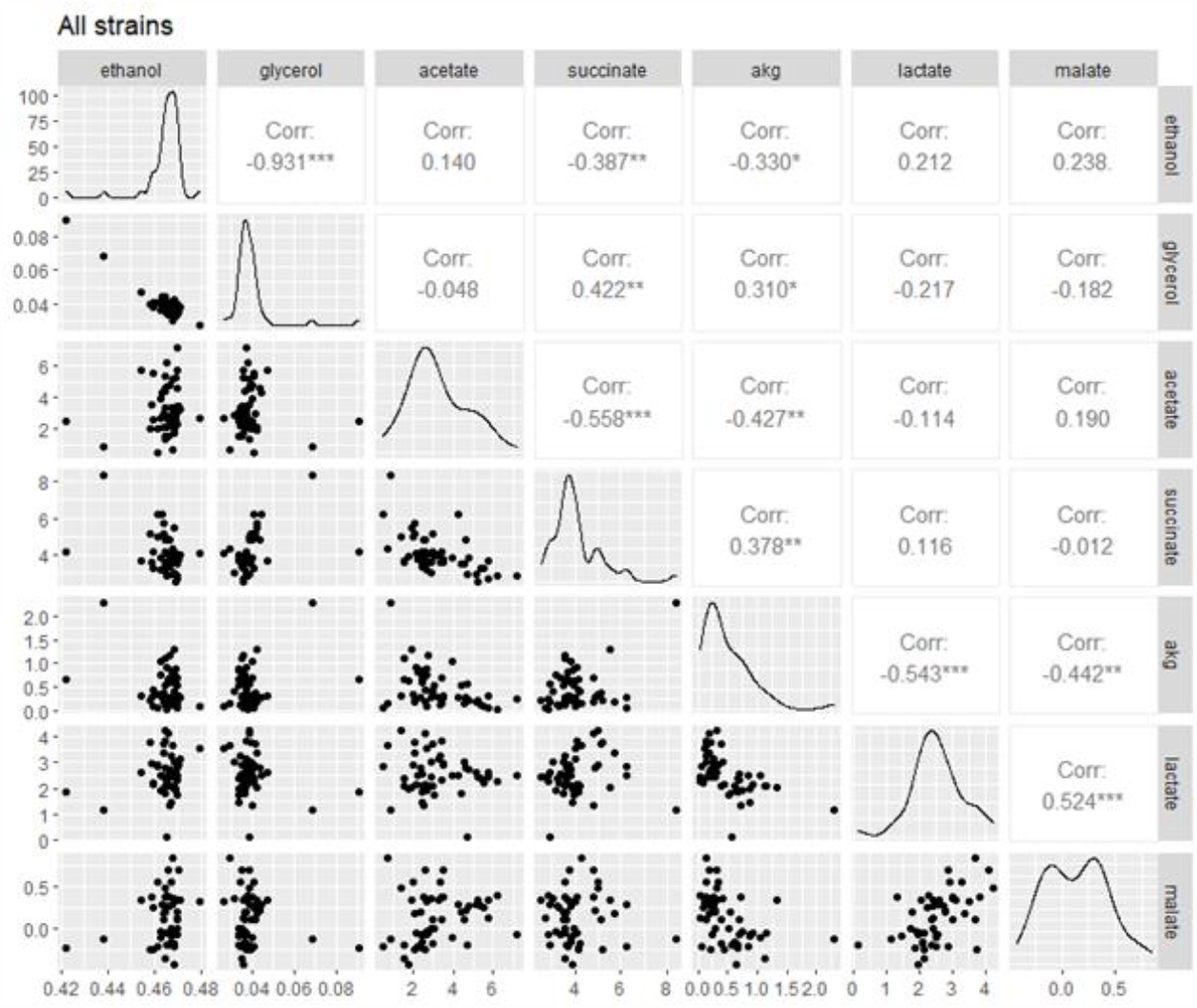
Pearson’s correlation matrix between all metabolic data for the whole set of strains (n = 51)

**Figure 3.**
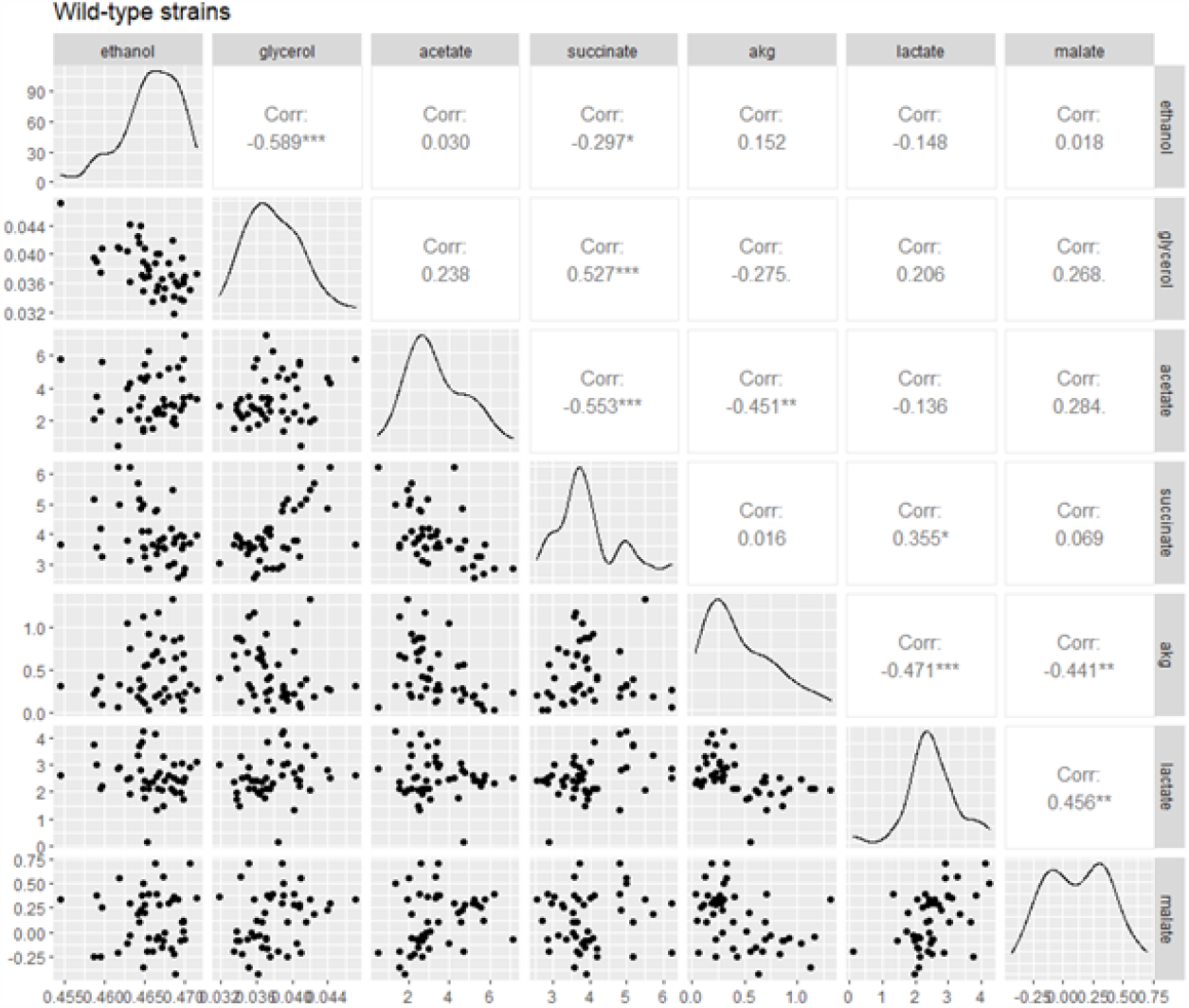
Pearson’s correlation matrix between all metabolic data for only wild-type strains (n = 46)

The strongest correlation spotted in the whole data set is a negative relation between glycerol and ethanol yields (r_all_ = -0.931) (figure 2). However, this correlation seems driven by modified and evolved strains because of their extreme yield values, as it is still present but weaker in the wild-type only data set (r_wild-type_ = -0.589) (figure 3). Other significant correlations (|r| > 0.300) were identified in both sets for other metabolites. This includes positive correlations between glycerol and succinate yields, lactate yield and malate difference, glycerol and α-KG yields and also negative correlations between acetate and succinate yields, acetate and α-KG yields, lactate and α-KG yields and α-KG yield and malate difference.

Most of the observed correlations apply to both the whole data set and the subset of wild-type strains. However, there are a few notable exceptions. In particular, two negative correlations, both involving ethanol, are significant in the whole data set, but are not apparent in the subset of wild-type strains. These negative correlations involve ethanol and succinate yields, as well as ethanol and α-KG yields. In addition, there is a positive correlation between α-KG and succinate yield that is only significant in the whole data set. Conversely, a positive correlation between succinate and lactate yield stands out in the wild-type subset but is not evident when the optimised and evolved strains are considered:. These distinctions between the two subsets can be directly attributed to the specific modifications and adaptations present in the optimised and evolved strains.

### Global analysis and hierarchical clustering

To obtain an overall view of our data set, a Principal Component Analysis (PCA) was performed with yields values for key metabolites: ethanol, glycerol, acetate, succinate, lactate and α-ketoglutarate (α-KG) and with malate which corresponds to the difference with the initial concentration (Figure 4). This analysis allowed us to position strains in relation to one another while providing insights on the impact of their genetic background. PCA was performed using wild-type strains only, to avoid biases induced by genetically modified and evolved strains. Hierarchical Clustering on Principal Components (HCPC) was also carried out on these results, allowing us to define three clusters of strains (Figure 4). This number of clusters was chosen because it is the smallest that best represents the distribution.

**Figure 4.**
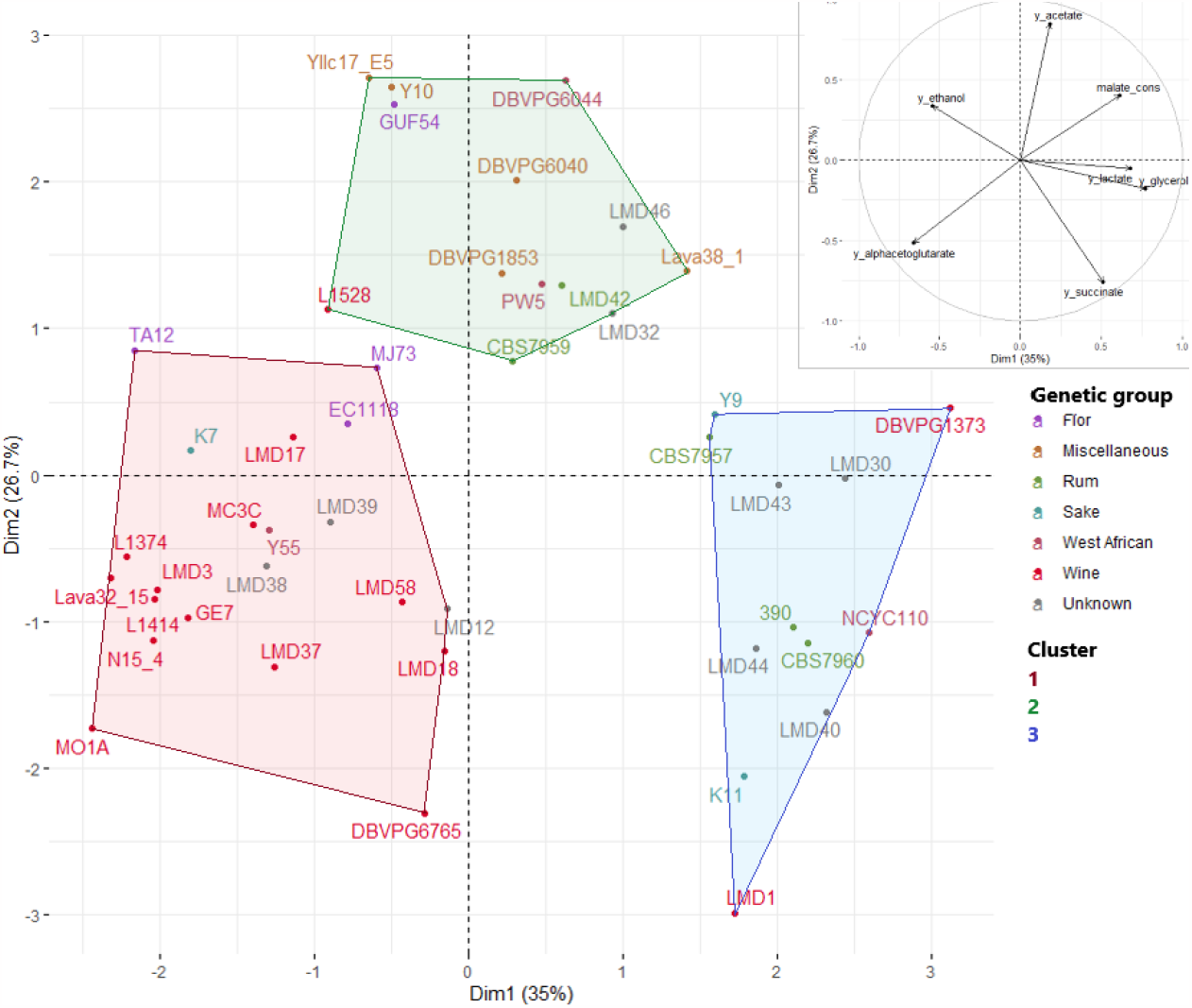
Representation of PCA on wild-type strains for ethanol, glycerol, acetate, succinate, α-KG and lactate yields and malate final difference with initial concentration, individual and variable plot, with HCPC clustering (Dimension 1 and 2, 61.7% of variance explained). *Coloured points represent strain, tinted by genetic origin. Three clusters were defined as: 1 (red), 2 (green) and 3 (blue)*

The wine strains group was quite homogenous and mainly located in cluster 1: 13 out of 16 wine strains are included. Moreover, strains LMD12, LMD38 and LMD39, which are commercialised for wine fermentation, are clustered with wine genetic strains. L1528, located in cluster 2, was still relatively close to cluster 1 in representation of dimension 1 and 2 (61.7% of variance explained). The wine group seems mainly driven by the high α-KG and low acetate yields, with high malate differences (except for the strain DBVPG1373). Most of flor strains (3 out of 4), including EC1118, were also located in cluster 1. Sake, rum and West African groups did not display any consistency in clustering and were scattered among clusters 2 and 3. However, rum strains, and unsequenced strains used in distillery conditions (LMD40, LMD43, LMD44 and LMD46) were relatively close in the representation, mainly driven by lactate and glycerol yields.

Finally, previously observed correlations between metabolites were still apparent and clear drivers of the clustering, such as the strong negative relation between α-KG yield and final malate difference.

### 3.4. Comparison with other works

Data obtained with the present screening strategy were compared with others arising from similar screening works (Camarasa *et al*., 2011; Legras *et al*., 2018; Nidelet *et al*., 2016). Data were normalised to consider only the relative distribution among a set. With this aim, the present work was included twice: first with all the strains and secondly with normalisation performed after withdrawing the modified strains (Figure 5).

**Figure 5.**
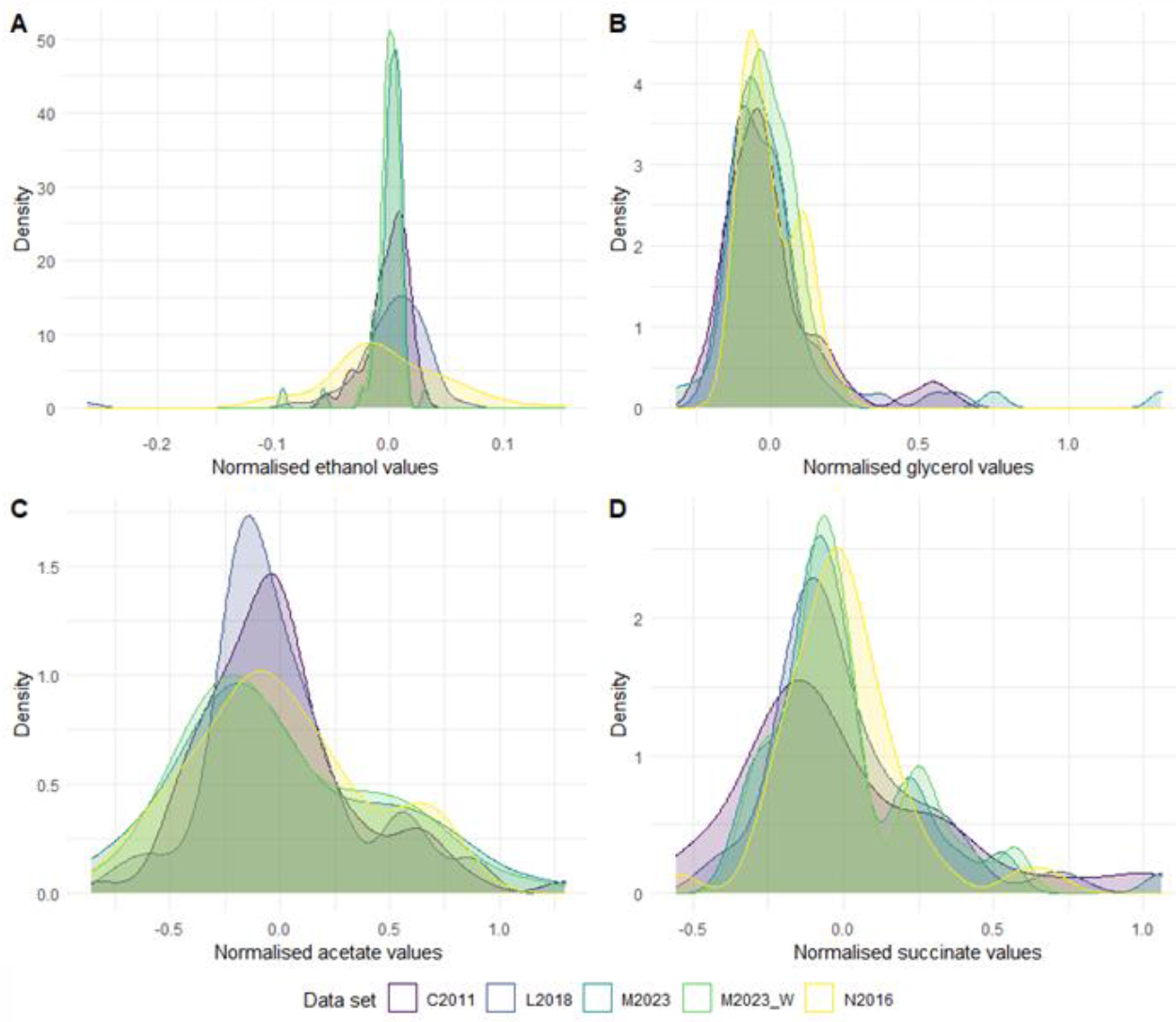
Distribution of ethanol (**A**), glycerol (**B**), acetate (**C**) and succinate (**D**) yields in 4 data sets. ***C2011***: *Camarasa* et al. *(2011);* ***N2016***: *Nidelet* et al. *(2016);* ***L2018***: *Legras* et al. *(2018);* ***M2023***: *this work, all strains;* ***M2023_W***: *this work, wild-type strains only*

All four metabolites showed a similar distribution, with ethanol always the less diversified yield and succinate and acetate displaying the most diversity. Glycerol showed an intermediary distribution. For all metabolites, divergent strains could be observed.

For ethanol, distribution seemed more limited in the data set exposed in this work, compared to the data set from the work of Nidelet *et al*. (2016), the two others being intermediary. Interestingly, this hierarchisation can be linked to the sampling time: the earlier the sampling, the higher the ethanol yield diversity among all strains.

## Discussion

*Saccharomyces cerevisiae* has already been the subject of multiple studies on its metabolism, with comparison between different strains and links between phenotype and genotype unveiled. Accordingly, this work on a diverse set of strains allows a broad view of primary metabolic diversity. Our results confirm significant variations that exist among different *S. cerevisiae* strains in terms of primary metabolites yields. The diversity of these variations is not uniform and depends on the specific metabolite considered. In this work, multiple correlations were confirmed (such as the well-known glycerol and ethanol negative correlation). Other correlations, between minor and less studied metabolites, were established as well, such as the positive link between glycerol and succinate, lactate and malate difference, glycerol and α-KG or the negative correlation between acetate and succinate, and between α-KG, lactate and malate difference. If most of the correlations are significant with or without evolved and genetically modified strains, slight differences appear between these two data sets. Moreover, this methodology affords a greater precision in metabolic yield assessment, corroborating existing data obtained in the last years and generating a robust and standardised dataset that can be reused in other studies on yeast metabolism. It enables yields to be precisely defined in a wine-like context, using a synthetic grape must with metabolite assessment at fermentation final stage. Lastly, it provides for the first time a screening of *S. cerevisiae* strains on lactate production in a wine-like context.

To simplify the discussion on metabolic yield results and their connections, a map of carbon central metabolism in oenological conditions is available in Figure 6.

**Figure 6.**
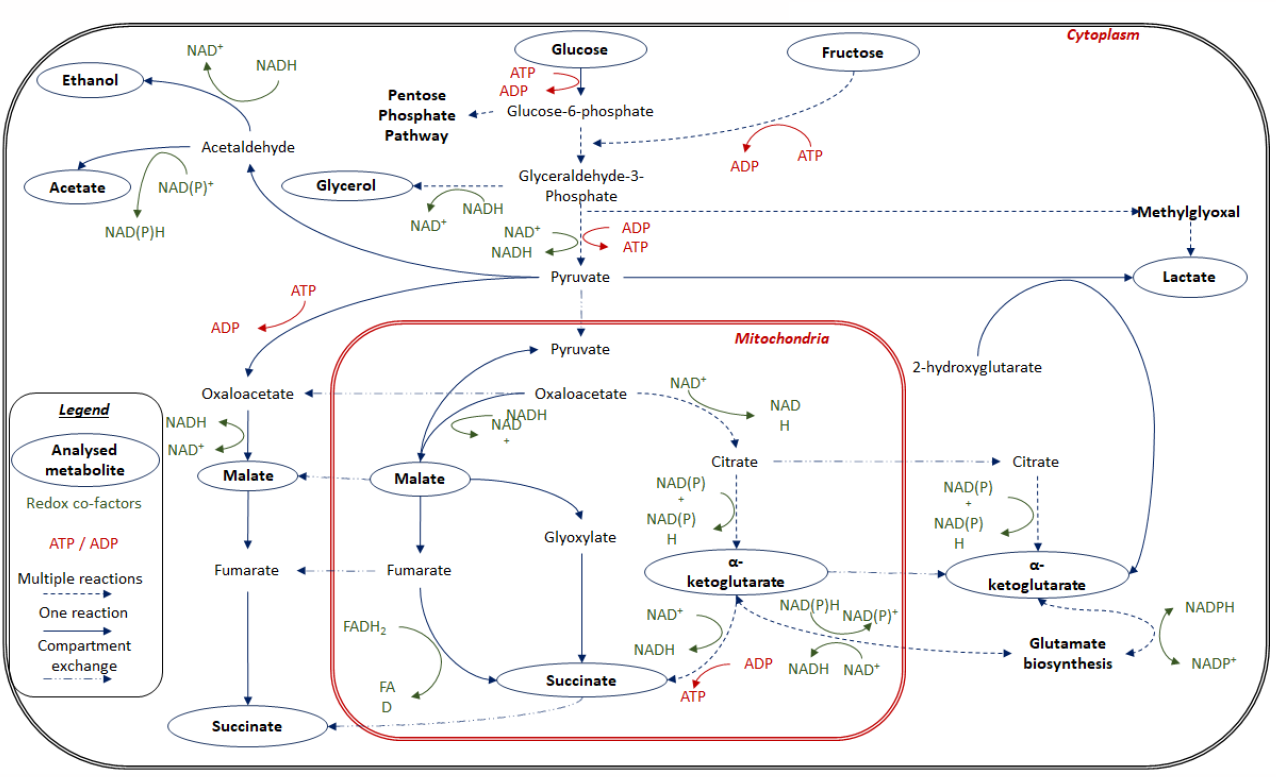
Simplified carbon central metabolism of *S. cerevisiae* in oenological conditions

The data set gathers 51 strains, including five evolved or modified strains for specific features linked to carbon metabolism. Although these strains were discarded for overall analysis, their behaviour in terms of metabolism was considered as control. We observed that the two strains displaying the highest glycerol yield and the lowest ethanol yield among all strains were LMD14 and 5074, both obtained following an adaptive evolution aiming to reduce their ethanol production while enhancing glycerol production (Tilloy *et al*., 2014). At the other end of the spectrum, LMD41, modified to enhance ethanol production while impeding glycerol production, exhibited the highest value of ethanol yield and the lowest for glycerol. Finally, the last GM strain LMD45 showed the second lowest acetate and glycerol yields, which is consistent with its modifications aiming to reduce fermentation by-product synthesis. All these features clearly correspond to the already known characteristics of the selected strains, for which they have been modified or evolved, and thus validate our methodology. The medium used in this study is a very close imitation of grape must, perfectly suited to study wine strain metabolism, but also suitable to every strain able to ferment a complex medium with high sugar concentrations (Bely *et al*., 1990). Fermentation duration being dependent on nitrogen level and temperature, our fermentations were carried out at 28°C with a must containing a relatively low concentration of sugars (from a winemaking point of view) and a high concentration of usually limiting nutrients (assimilable nitrogen, vitamins, anaerobic growth factors…). This ensured a rapid and complete hexose conversion to ethanol (Rollero *et al*., 2015).

In overview, if we compare metabolites with one another, great differences in yield exist. Ethanol was the most produced compound, with a yield ten-fold higher than glycerol, whose yield was itself ten-fold higher than acetate yield. α-KG had the lowest yield values but still close to acetate. It should be noted that substantial yield variations were observed among strains for each metabolite, underlining the suitability of our experimental conditions to effectively discriminate strains on the basis of their primary metabolite yields.

However, variations among yields differed depending on the considered metabolite. Ethanol and glycerol are the most produced metabolites during alcoholic fermentation. Here, ethanol, with a variation coefficient inferior to 2%, showed a remarkably low variation. This level of variation was even lower when considering only wild-type strains. By contrast, glycerol yield exhibited a higher degree of variability, with a coefficient of variation around 25%. The same variation ranking can be observed in Nidelet *et al*. (2016) results, obtained in a similar medium using 43 strains (including 20 in common with our set), with ethanol being the most constant flux, followed by glycerol and then acetate, succinate and α-KG as the most variable. For the first four metabolites, these observations were confirmed by our normalised comparison and can also be found in other different works on CCM: indeed, Tronchoni *et al*. (2022), performed a screening in wine-like media in aerobic conditions using 25 *S. cerevisiae* wine strains. Ethanol yields were lower than in our results, which is consistent with aerobic conditions, but the range of variation was very similar: no major observable differences and significant differences only between strains with extreme values. Another comparable screening can be found in the work of Nieuwoudt *et al*. (2006) on 15 strains (commercial or not) and 19 hybrids using both natural and synthetic laboratory must. On both media, similar results were obtained: a higher range of variation was observable for glycerol than for ethanol. Similarly, Hubmann, Foulquié-Moreno, *et al*. (2013) performed a relevant screening on 52 beer and distillery strains of *S. cerevisiae* for their ethanol and glycerol yields, on a YPD-like medium. All these data present a greater diversity among strains for glycerol than for ethanol.

Furthermore, our data confirm the negative correlation observed in prior studies between ethanol and glycerol productions or yields, with a stronger link when extreme values from modified and evolved strains (strains 5074, LMD14 and LMD41) are considered. These values clearly drive the correlation in the whole set, but this negative correlation is still significant in the wild-type strains set, and demonstrate the results of redox balancing between the two metabolites (Goold *et al*., 2017). Glycerol, considering its concentration and variation range and the strong negative correlation with ethanol yield, is confirmed to be the best candidate to orient carbon fluxes in the cell toward ethanol production. Surprisingly, no relation between belonging to a genetic group and yield of glycerol or ethanol was found in our data. This goes against previous observations stating that wine strains are high glycerol producers compared to other groups (Camarasa *et al*., 2011). Nevertheless, it is worth noting that, in the study of Camarasa *et al*. (2011), groups were based on their environmental origin whereas ours were based on genetic origin. However, these two origins do not always match (as an example, strain Y55 used to be classified as a laboratory strain isolated from a wine environment, but Liti *et al*. (2009) showed that this strain is in fact closer to a West African genetic lineage).

Acetate is responsible for major off-flavours in wine, and so is subjected to regulatory limits (Paraggio and Fiore, 2004; Vilela-Moura *et al*., 2008). Our data revealed substantial variation in acetate yield and significant links with strains from different genetic groups. Specifically, strains from the wine group displayed a very low acetate yield, significantly lower than those from the West African group. This phenotype can be seen as a domestication footprint as it is most likely a direct consequence of selection for low acetate production during wine fermentation, while West African strains are known to have been less subjected to domestication (Warringer *et al*., 2011).

Succinate was produced in minor concentrations compared to ethanol or glycerol and its production appears correlated negatively with ethanol and consistently positively with glycerol, with whom it shares a similar variance (all strains considered). One explanation to this observation could be linked to the redox balance (Figure 6). Indeed, NADH used in glycerol production needs to be regenerated afterwards. In the meantime, production of succinate by the oxidative branch of TCA produces NADH while this branch is also the most subject to variation (Camarasa *et al*. 2003) (Figure 6).

Another metabolite, α-KG, provided interesting results, especially with its link to specific genetic groups. Wine strains showed a higher yield than other groups, while rum strains showed a lower yield. This particularity can be directly explained by the strong relation between this metabolite and nitrogen metabolism. Indeed, α-KG is mainly used in the cell for ammonium uptake and subsequent glutamate synthesis (Figure 6). However, in grape must (and also in the synthetic must used in this study), where glutamate is abundant, α-KG is not fully utilised and is instead released into the environment (Avendaño *et al*., 1997; DeLuna *et al*., 2001; Camarasa *et al*., 2003; Magyar *et al*., 2014). Glutamate synthesis consumes the NADPH cofactor, which must be regenerated. One way to produce NADPH is the conversion of acetaldehyde to acetate from NADP^+^ (Saint-Prix *et al*., 2004) (Figure 6). Moreover, the wine strain group displayed a low acetate yield on average (if we exclude DBVPG1373 which shows abnormal values compared to the rest of the group). In the work of Nidelet *et al*. (2016), it was observed that acetate flux in fermentation is negatively correlated to biomass synthesis, itself negatively correlated to α-KG. Despite the lack of data on biomass, the negative link between acetate and α-KG on our set confirms these results. Moreover, succinate and α-KG were negatively correlated in the whole data set, but not when considering only wild-type strains. This correlation was mainly driven by strain LMD14, which has been developed using adaptive evolution for high glycerol and low ethanol production (Tilloy *et al*. 2014). This strain is also known for its high organic acid production, here confirmed and identified as a consequence of a high TCA activity (Camarasa *et al*., 2003).

Another organic acid of interest in a wine context is lactate, which has not been extensively studied in *S. cerevisiae*, and is produced in minor quantities from malate, mainly as a result of bacterial activity (Volschenk *et al*., 2006). Yet, lactate production in *S. cerevisiae* displayed a wide range of diversity, with significant links with genetic groups. Wine and flor strains tended to be low producers of lactate whilst rum and sake strains were high producers. Additionally, all strains ungrouped but used in distillery conditions (LMD40, LMD44, and GM strains LMD45 and LMD41) ranked among the best producers (available in supplementary information). These findings enforce the idea of a connection between the rum group and lactate production. However, if diversity was indeed present, measured concentrations were by far lower than those observed in wine fermentation with other yeasts such as *Lachancea thermotolerans* that can reach up to 12 g/L (Vicente *et al*. 2021). Contrary to *Lachancea* yeasts, which harbour lactate dehydrogenase dedicated enzymes (LDH), lactate in *S. cerevisiae* is produced in small amounts from pyruvate reduction by residual lactate dehydrogenases (coded by *DLD1, DLD2* and *DLD3* genes), Dld3p being responsible for the major part of LDH activity in anaerobic conditions (Figure 6). This activity is coupled to α-KG production from 2-hydroxyglutarate and pyruvate (Becker-Kettern *et al*., 2016). The present data set showed a negative correlation between lactate and α-KG. This can be explained by the multiple paths available leading to α-KG synthesis, especially in nitrogen metabolism, but also by a second way for lactate production in yeast. Indeed, lactate is also produced during methylglyoxal degradation. This toxic metabolite is produced from glycolysis intermediary (trioses phosphates) and can be degraded through the glyoxalase pathway. Moreover, our fermentation conditions, with quite high hexose concentrations, entailed a high glycolytic flux that can cause a harmful methylglyoxal production harmful to cells (Martins *et al*., 2001; Stewart *et al*., 2013).

Finally, the last organic acid, malate, stands in a particular situation in wine fermentation: it was already present in relatively high concentration in the synthetic grape must used (6 g/L) and it can be both consumed and produced during fermentation. Therefore, its analysis as a final concentration gives only an indication on the final balance resulting from the whole metabolic changes (Vion *et al*., 2023). This led to the identification of strains resulting in malate production (mostly rum and west African strains), and others, mainly wine strains, displaying no or little malate consumption. This result is counterintuitive since malate is naturally present in the natural environment of wine strains. In the whole set, malate consumption never exceeded 1 g/l (see supplementary information), which is consistent with previous results obtained in a large scale study on *S. cerevisiae* (Yéramian *et al*. 2007). Final malate concentration is found to be negatively correlated to α-KG content, meaning that higher malate production or lower consumption was associated with lower α-KG production. Without making further hypotheses, these results point out the differences in TCA cycle activity between strains and between genetic groups, making it a metabolic pathway of carbon metabolism more diverse than ethanol or glycerol production. Overall, tendencies in our data set are consistent with conclusions drawn in other publications, including those used to select our set of strains (Camarasa *et al*., 2011; Nidelet *et al*., 2016, Legras *et al*., 2018). Our work thus confirms crucial observations on metabolism and comforts prior information. Strains from the West African genetic group (which groups strains from palm wine and other traditional African beverages making processes) and from the flor group have been identified as very high acetate and low succinate producers. Acetate showed a great diversity, larger than glycerol or ethanol, in our set, as already shown by Tronchoni *et al*. (2022).

Moreover, the individual metabolite approach highlighted interesting correlations. PCA also allowed to visualise all these correlations combined, which makes it a good tool to group strains according to their metabolic features. For wine strains, metabolic clusters on the PCA matched with the genetic groups except for a few strains. Rum strains and unsequenced strains used in distillery conditions (LMD40, LMD43, LMD44 and LMD46) were grouped in cluster 2 and 3 by Hierarchical Clustering on Principal Components, but relatively close in dimension 1 and 2 of PCA representation, suggesting a phenotypic proximity linked to the environment. Genome sequencing would be necessary to conclude on the place of the four commercial strains in the rum group. The 3 strains from the sake group were scattered in the PCA representation, without any common features appearing. This high phenotypic diversity has already been highlighted in the past for other traits in the sake population (Warringer *et al*., 2011).

The ability to complete a wine-like fermentation is strongly linked to domestication and genetic origin (strains from bread or from natural environments such as oak trees are most of the time unable to perform a wine-like alcoholic fermentation (Camarasa *et al*., 2011; Legras *et al*., 2018; Tapia *et al*., 2018)); yet, a complementary set of strains, wider and more balanced between genetic groups, could provide more diversity and strengthen our analysis of natural yield variations. Strains from other anthropic origins (such as beer or cider) were not included here. However, if it appears that these strains can complete a full alcoholic fermentation in our conditions, they should be considered in an extended study. Moreover, for strains from genetic groups other than wine, a synthetic grape must can represent conditions very far from their usual environment. However, despite these differences that can be overcome with scale adjustment, our methodology provides keys to identify strains with good potentialities for wine fermentation.

Multiple studies have compared numerous strains for their primary metabolites production in fermentation. However, our study compares strains from different genetic groups on a wine-like medium, only focusing on complete fermentation concentrations. Here we confirm precedent observations but also provide a robust comparative methodology and an easily usable data set obtained on 51 strains from various genetic backgrounds. Experimental conditions allowed a medium-throughput screening, which is a good balance between phenotyping a large number of strains and having a high accuracy enabling the identification of traits with low variation. This screening helps to define and confirm the existing phenotypic variations for wine fermentation products among the *S. cerevisiae* species and sets the potential of improvement for these traits. It also provides information on lactate production in *Saccharomyces cerevisiae*, which shows a poor ability, however with a significant diversity and links with genetic origins. Nevertheless, the set of primary metabolites considered here is limited, without any data on notable aromatic metabolites, positively or negatively impacting wine fermentation (acetaldehyde, esters, higher alcohols, acetoin…). Moreover, these other phenotypes have been shown to exhibit a greater diversity among *S. cerevisiae* strains than primary metabolites, even in homogenous genetic groups. Completing this analysis with additional information on other metabolite production or consumption would strengthen the clustering and allow a broader view on metabolism differences between strains while revealing patterns of interaction between pathways (nitrogen metabolism, lipid biogenesis…). Finally, this would reveal precisely strain relevance for further development projects, applied to wine or other fields.

To conclude, the present screening answers the main initial questions: some diversity, weak but significant, exists in ethanol yield among the *S. cerevisiae* species. Larger fluxes, such as ethanol or glycerol, are the most constraint and not linked to genetic origins, while, by contrast, smaller fluxes show larger variations and clear links with genetic origin. This represents improvement potentials of wine strains for these characteristics with non-GM methods (such as adaptive laboratory evolution, positive selection, breeding…). If the two major produced metabolites, ethanol and glycerol, are linked in their production, the yield of minor metabolites is more related to the genetic background of strains which is shaped by selection in a defined environment. Beyond confirming results observed in the last years with a robust and standardised method, our work also provides insights on little-studied metabolites with high technological potential in wine fermentation.

## Abbreviations

CCM: Carbon Central Metabolism;
GM: Genetically Modified;
CV: Coefficient of Variation;
α−KG: α-Ketoglutarate;
TCA: Tricarboxylic Acid;
YAN: Yeast Assimilable Nitrogen

## Acknowledgements

The authors would like to thank Jean-Luc Legras and Carole Camarasa for helpful discussions. They also thank Philippe Chatelet for useful suggestions on the text. The authors thank the University of Azores, Ricardo Franco-Duarte, Marie-José Ayoub, as well as the National Research Institute of Brewing (Hiroshima, Japan) and all the international yeast collections for providing strains. Finally, the authors thank Anne Ortiz-Julien from Lallemand SAS for her support of this work.

## Data, scripts, code, and supplementary information availability

Data sets, R scripts and all supplementary informations are available on the following link: **10.5281/zenodo.8425971**

